# Nanoscale Structure Determination of Murray Valley Encephalitis and Powassan Virus Non-coding RNAs

**DOI:** 10.1101/2020.01.10.901082

**Authors:** Tyler Mrozowich, Amy Henrickson, Borries Demeler, Trushar R Patel

**Author notes:** Corresponding Author: Trushar R. Patel.

## Abstract

Viral infections are responsible for numerous deaths worldwide. Flaviviruses, which contain RNA as their genetic material, are one of the most pathogenic families of viruses. There is an increasing amount of evidence suggesting that their 5’ and 3’ non-coding terminal regions are critical for their survival. In this study, the 5’ and 3’ terminal regions of Murray Valley Encephalitis and Powassan virus were examined using biophysical and computational modeling methods. First, the purity of *in-vitro* transcribed RNAs were investigated using size exclusion chromatography and analytical ultracentrifuge methods. Next, we employed small-angle X-ray scattering techniques to study solution conformation and low-resolution structures of these RNAs, which suggested that the 3’ terminal regions are highly extended, compared to the 5’ terminal regions for both viruses. Using computational modeling tools, we reconstructed 3-dimensional structures of each RNA fragment and compared them with derived small-angle X-ray scattering low-resolution structures. This approach allowed us to further reinforce that the 5’ terminal regions adopt more dynamic structures compared to the mainly double-stranded structures of the 3’ terminal regions.

## 1. Introduction

Family Flaviviridae are small, positive-sense single-stranded RNA viruses that replicate within host cells of arthropods and/or vertebrates. Flaviviruses include deadly viruses such as Murray Valley Encephalitis Virus (MVEV), Powassan Virus (PowV), Japanese encephalitis virus, Dengue virus, Zika virus, West Nile virus and Yellow Fever virus. The World Health Organization and the Centers for Disease Control both cite flaviviruses as a global health threat owing to the ease of transmission by mosquitoes and lack efficient therapeutic or immunoprophylactic strategies [1]. Given the magnitude and severity of disease from this class of viruses, there is a critical need for therapeutics; however, the limited understanding of the virus’s replication and their complex interactions with the host cellular proteins through terminal region (TR) interactions hinders therapeutic development [2-4].

MVEV is a member of the Japanese encephalitis serological complex of flaviviruses and was first isolated in 1951 during the initial outbreak in Australia [5]. It is believed to be maintained at the top end of the Northern Territory and the North of Western Australia by a cycle involving mosquitoes (*Culex annulirostris*) and birds [6]. The last major outbreak was in 1974 where 58 cases were reported, 20% resulted in death [7]. Since 2011 there have been increasing instances of MVEV, including nine cases in 2011, three of which resulted in death [8,9]. PowV is the only member of the tick-borne encephalitis serogroup that is currently present in North America [10]. PowV has seen a drastic increase in the last 18 years, compared to the previous 40 with an increase of 671% [11]. It was first identified in Powassan, Ontario in 1958 [12] and has become endemic in the upper Midwest and the Northeastern United States, but cases have also been reported in eastern Russia [13]. The increase in PowV incidences over the last decade highlights the importance of conducting further research to understand the structure-function relationships of the viral genome to ultimately develop therapeutic approaches to combat such viral infections.

Flaviviral genomes consist of a single-stranded RNA molecule approximately 10-11k nucleotides long, depending on species. The RNA genome contains one single open reading frame (ORF) which is flanked by terminal regions (TRs) [1]. The ORF codes for a single polypeptide that is cleaved by a combination of viral and host proteases to produce three structural proteins and the seven non-structural proteins [14]. The flanking structural TRs are highly conserved across all flaviviruses and appear as complex folding sequences essential for viral replication [1,15-17]. Since these terminal regions are so highly conserved and intolerable to mutations/deletions, it is hypothesized, and subsequently shown, that these regions interact with the specific host proteins required for replication [18-20]. The important role of these terminal regions is also highlighted by their ability to cyclize, where the 3’ TR interacts with the 5’ TR [21,22]. This cyclization of the viral genome is essential for the correct positioning of the NS5 protein, produced by the viral genome [23], which is the viral RNA dependent RNA polymerase, which is required to produce (-) sense stranded RNA, which is used as the replication template for further production of the viral (+) sense RNA genome, facilitating viral replication [24]. Furthermore, flaviviral TRs also interact with a variety of host proteins [21,25,26]. Thus, it is essential to study the complex structure of the TRs to further understand TRs-viral/host protein interactions and cyclization events, as they play an essential role in viral replication.

Although several attempts have been made to study RNA structures in general [27,28], detailed insight into the structural features of multiple flaviviral strains is lacking. Such detailed insight could provide a platform to develop novel antiviral therapies as well as further our knowledge about viral replication events. Therefore, using a combination of biophysical techniques and computational calculations, we have constructed 3-dimensional structures for the 5’ and 3’ TRs of both MVEV and PowV. This study provides insight into the organization of flaviviral TRs and provides a framework for the characterization of other flaviviral TRs, or TR segments or other non-coding RNAs.

## 2. Materials and Methods

### 2.1.1 RNA Preparation and Purification

We prepared cDNA sequences under the control of a T7 RNA polymerase promoter with two additional G nucleotides on the 5’ end, followed by an XbaI restriction enzyme cut site (T^CTAGA). We designed our constructs for MVEV and PowV based on the Genebank sequences of KX229766.1and EU670438.1, respectively. All RNA constructs used in the experiments are listed as follows:

1, MVEV 5TR 1-96nt GGAGACGUUCAUCUGCGUGAGCUUCCGAUCUCAGUAUUGUUUGGAAGGAUCAUUGAUUAACGCGGUUUGAACAGUUUUUUGGAGCUUUUGAUUUCAAU

2, MVEV 3TR 10914-11014 GGCCUGGGAAAAGACUAGGAGAUCUUCUGCUCUAUUCCAACAUCAGUCACAAGGCACCGAGCGCCGAACACUGUGACUGAUGGGGGAGAAGACCACAGGAUCUU

3, POWV 5TR 1-111 GGAGAUUUUCUUGCACGUGUGUGCGGGUGCUUUAGUCAGUGUCCGCAGCGUUCUGUUGAACGUGAGUGUGUUGAGAAAAAGACAGCUUAGGAGAACAAGAGCUGGGAGUGGUUU

4, POWV 3TR 10735-10839 GGCCCCCAGGAAACUGGGGGGGCGGUUCUUGUUCUCCCUGAGCCACCACCAUCCAGGCACAGAUAGCCUGACAAGGAGAUGGUGUGUGACUCGGAAAAACACCCGCUU

Each RNA construct was *in-vitro* transcribed using T7 RNA polymerase then purified using a Superdex 200 increase (GE Healthcare) on an ÄKTA pure FPLC system (GE Healthcare) at 0.5mL/min. RNA Fractions were collected and pooled from the size exclusion chromatography (SEC) fraction collector. Pooled fractions were ethanol precipitated and resuspended in RNA buffer (50 mM Tris pH 7.5, 100 mM NaCl and 5 mM MgCl_2_). The pooled fractions were analyzed using Urea-Poly Acrylamide Gel Electrophoresis (Urea-PAGE). 10 µL of ∼500 nM RNA was mixed with 2 µL of RNA loading dye and loaded into a 1.0 cm well PAGE casting plate (BioRad). Urea-PAGE (11.2%) was run at 300V at room temperature for 20 minutes in 0.5 X TBE, followed by staining with Sybr Safe (Thermofisher Scientific) and UV visualization. Prior to any experiment (AUC or SAXS) RNA was heated to 95 ºC for 5 min and cooled passively to room temperature to facilitate refolding.

### 2.2 Analytical Ultracentrifugation (AUC)

The sedimentation velocity (SV) – AUC data for FPLC-purified RNA samples were collected using a Beckman Optima centrifuge and AN50-Ti rotor at 20 °C. Samples were loaded into Epon-2 channel centerpieces at 0.6 µM, 0.5 µM, 0.6 µM and 0.58 µM for 5’ and 3’ TRs of MVEV and PowV, respectively in 50 mM Tris pH 7.5, 100 mM NaCl and 5 mM MgCl_2_ buffer and centrifuged at 35,000 revolutions per minute. Scans were collected at every 20 seconds. We used the UltraScan-III package [29] and San Diego Supercomputing Center on Comet as well as the Texas Advanced Computing Center on Lonestar-5 computing facility for data processing. The SV-AUC data were initially analyzed by means of two-dimensional spectrum analysis (2DSA) with simultaneous removal of time-invariant noise, the meniscus and position fitted [30], enhanced van Holde-Weischet protocol [31], genetic algorithm refinement [32] and further refined with Monte Carlo analysis [33]. The density and viscosity corrections for the buffer were estimated with UltraScan to be 1.0030g/cm3 and 1.0100 cP.

### 2.3 Small angle X-ray scattering (SAXS)

Utilizing the B21 beamline at Diamond Light Source (Didcot, UK), small-angle X-ray scattering (HPLC-SAXS) data was collected as previously described [34]. Making use of an in-line Agilent 1200 (Agilent Technologies, Stockport, UK) HPLC connected to a flow cell, each purified RNA, 50 µL of ∼2.0 mg mL^-1^ were injected into a buffer equilibrated Shodex KW403-4F (Showa Denko America Inc.) size exclusion column at a flow rate of 0.160mL min^-1^. Each frame was exposed to the X-rays for 3 s. Each sample peak region was integrated, using a combined total of approximately 10 datasets, buffer subtracted and then merged using Primus [35] or ScÅtter [36] as described previously [37]. The merged data were then analyzed using Guinier approximation to obtain the radius of gyration (*R*_*g*_) and study homogeneity of samples [38]. We also performed dimensionless Kratky analysis [39] to investigate if the RNA molecules of interest are folded, as reviewed earlier [40]. Next, the pair-distance distribution (*P*(*r*)) analysis was performed using the program GNOM [41], which provided the *R*_*g*_ and the maximum particle dimension (*D*_*max*_, the radius at which the *P*(*r*) dependence approaches to zero). Information from the *P*(*r*) plot was then used to generate the models using DAMMIN [42], with no enforced symmetry (P1), as described earlier [43]. The resulting models were averaged, and filtered to obtain a singular representative model using the DAMAVER package [44] as described previously [45,46].

### 2.4 Atomic Structures Calculations

MC-Fold [47] was used to predict alternative low energy secondary structures for 5’ and 3’ TRs of MVEV and PowV (Figure 1). We selected the lowest energy structures in each case as input files for MC-Sym that allows fragment-based reconstruction of 3-dimensional structures using known structures, as described earlier [47]. Briefly, 100 all-atom structures for each RNA were calculated, which were minimized using protocols implemented in MC-Sym. Scattering profiles were simulated with CRYSOL based on minimized structures and compared to experimental SAXS data…The minimized structures were subjected to CRYSOL to determine *R*_*g*_ and goodness-of-fit parameter (*χ*^*2*^) [48]. The MC-Sym derived structures were ranked based on their *χ*^*2*^ values and were aligned with low-resolution structures using program SUPCOMB [49].

**Figure 1.**
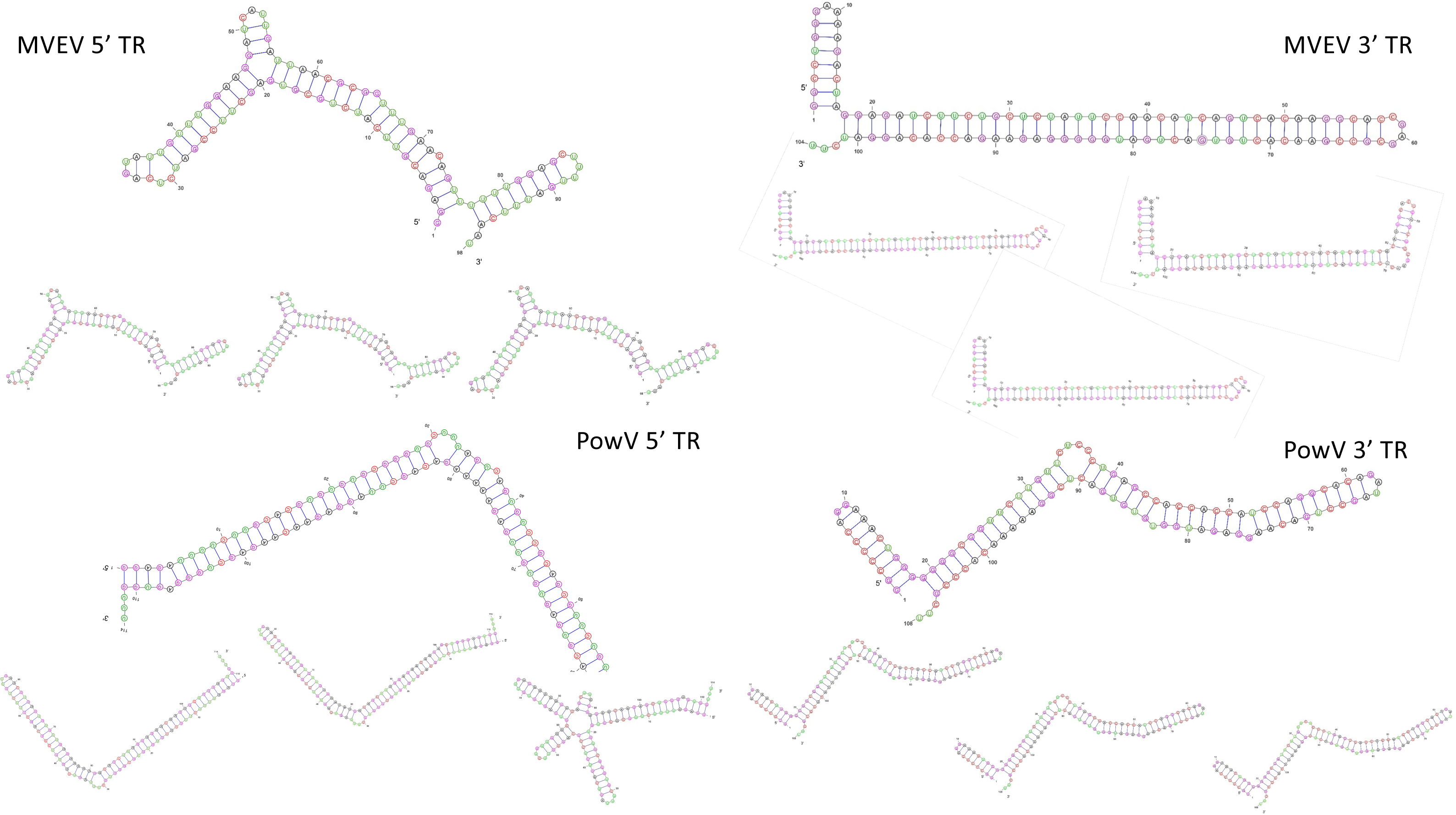
Lowest energy predicted secondary structures of MVEV and PowV non-coding RNA regions calculated using VARNA [69]. ΔG values for the lowest energy structures for MVEV 5’TR, MVEV 3’TR, PowV 5’TR, and PowV 3’TR are −86.41, −104.44, −103.07, and −103.07 kcalmol^-1^, respectively.

## 3. Results

### 3.1. Purification of in-vitro transcribed RNA

The *in-vitro transcribed* RNAs were purified using a Superdex 200 increase column connected to the ÄKTA FPLC unit. Figure 2A presents an elution profile for all four RNA constructs, where peaks at ∼8 mL represent elution of template plasmids used to *in-vitro* transcribe these RNAs. The RNAs of interest eluted at ∼13 to 14.5 mL and the peaks ∼11 mL suggest the presence of an oligomeric assembly of RNAs. The monodispersed fractions (at ∼13 to 14.5 mL) were pooled together and subsequently analyzed using Urea-PAGE (Figure 2B). As demonstrated in Figure 2B, all four RNAs migrate similarly and are highly pure, devoid of any aggregation or degradation. These fractions were stored at 4 C until further experiments were carried out.

**Figure 2.**
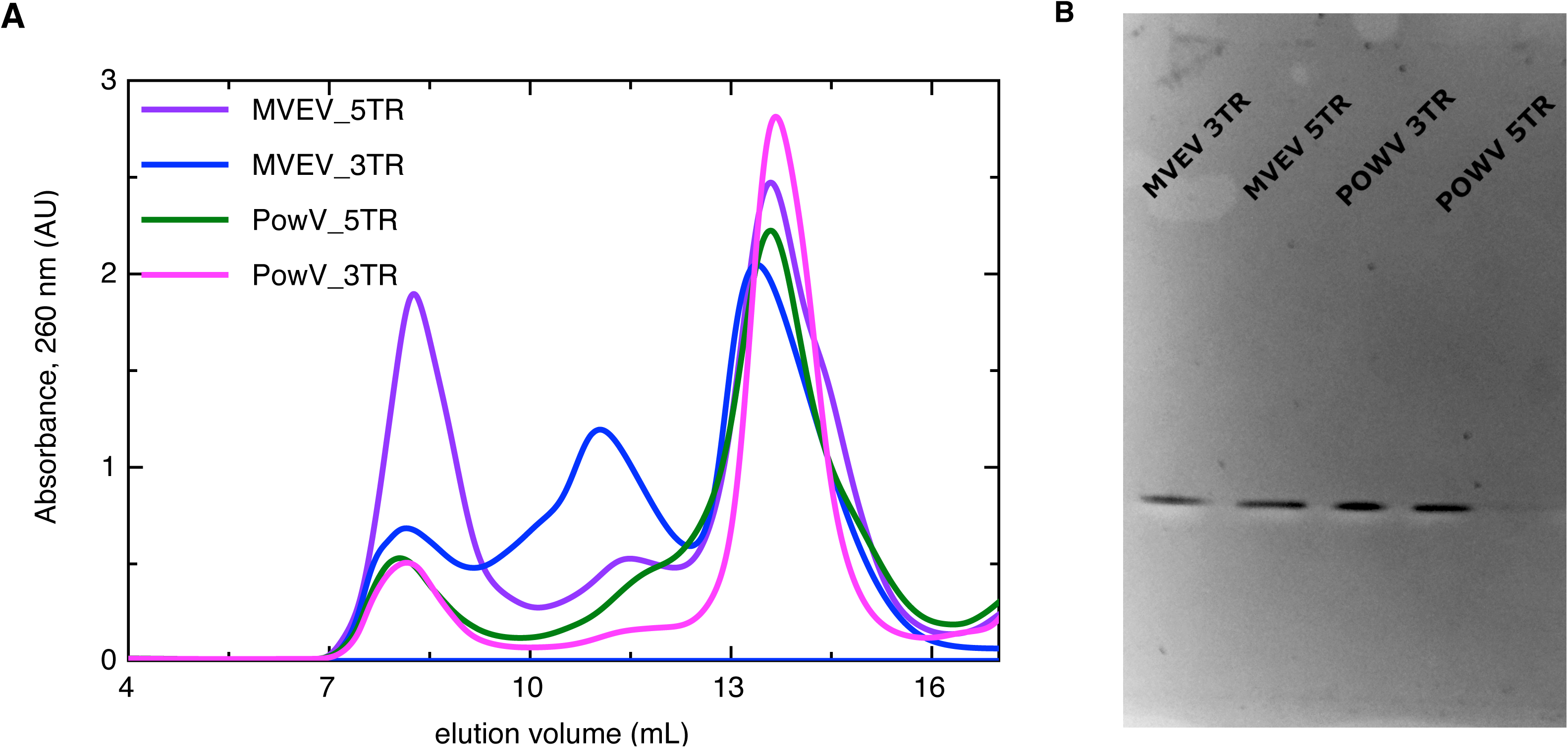
Purification of in-vitro transcribed 5’ and 3’ TRs of MVEV and PowV RNA. (**A**) SEC elution profile of purified RNAs where x-axis represents elution volume and y-axis represents absorbance at 260 nm; (**B**) Urea-PAGE demonstrating purity of the individual pooled RNA fractions from SEC peaks (∼13 to 14.5 mL).

### 3.2. Homogeneity studies of RNA

Analytical Ultracentrifuge is a versatile technique to study the homogeneity of biomolecules in solution [50]. In order to further evaluate the purity of all four purified RNAs, we performed sedimentation velocity experiments at the concentration range of 0.5-0.7 µM, and processed the data using UltraScan [29] as described in the Materials and Methods section. Figure 3 presents the sedimentation coefficient distribution for MVEV 5’ and 3’ as well as PowV 5’ and 3’ RNA fragments. The sedimentation velocity analysis suggests that all four RNAs are mainly monodisperse with sedimentation coefficient values of 4.27 S for MVEV 5’ TR, 4.30 S for MVEV 3’TR, 4.49 S for PowV 5’ TR and 4.53 S for PowV 3’ TR (S=10^−13^ seconds), as summarized in Table 1. Note that the peaks at ∼ 5.5 S suggest that all four RNAs form dimeric or high-order conformations in solution. The AUC data also suggest that PowV TRs with a *M*_*w*_ of ∼38 kDa have a slightly higher sedimentation coefficient compared to the MVEV TRs with *M*_*w*_ of ∼32 kDa (Table 1). Overall, these experiments indicate that all four RNAs are of suitable purity to perform HPLC-SAXS to determine solution structure.

**Table 1.** Biophysical parameters of MVEV and PowV non-coding RNAs.

| Sample | MVEV 5'TR | MVEV 3'TR | PowV 5'TR | PowV 3'TR |
| --- | --- | --- | --- | --- |
| Mw (kDa, sequence) | 31.38 | 31.57 | 36.92 | 37.80 |
| Sedimentation coefficient (S)<sup>▽</sup> | 4.27<br>(4.08, 4.46) | 4.30<br>(4.20, 4.41) | 4.50<br>(4.42, 4.55) | 4.53<br>(4.52, 4.53) |
| $I(0)^{\#}$ | $0.0098 \pm 4.7 \times 10^5$ | $0.019 \pm 1.4 \times 10^5$ | $0.019 \pm 6.7 \times 10^5$ | $0.034 \pm 8.2 \times 10^5$ |
| $q.R_g$ range | 0.38 – 1.26 | 0.59 – 1.26 | 0.34 – 1.26 | 0.47 – 1.29 |
| $R_g$ (Å)<sup>#</sup> | $29.01 \pm 0.22$ | $45.72 \pm 0.48$ | $34.65 \pm 0.19$ | $35.84 \pm 0.13$ |
| $I(0)^{\#\Delta}$ | $0.098 \pm 3.8 \times 10^5$ | $0.018 \pm 8.1 \times 10^5$ | $0.019 \pm 4.2 \times 10^5$ | $0.034 \pm 8.9 \times 10^5$ |
| $R_g$ (Å)<sup>Δ</sup> | $30.16 \pm 0.11$ | $46.16 \pm 0.18$ | $34.48 \pm 0.06$ | $38.39 \pm 0.08$ |
| $D_{max}$ (Å)<sup>Δ</sup> | 100 | 150 | 105 | 130 |
| $\chi^2^*$ | ~0.78 | ~0.83 | ~0.78 | ~0.80 |
| NSD<sup>*</sup> | 0.71 ± 0.01 | 0.80 ± 0.02 | 0.85 ± 0.02 | 0.72 ± 0.01 |

$\nabla$ - determined using SV-AUC analysis and UltraScan-III package [67]. Sedimentation coefficients obtained following genetic algorithm-Monte Carlo analysis. Measured value represents the mean. Bracketed values represent 95% confidence interval from Monte Carlo analysis.

**Figure 3.**
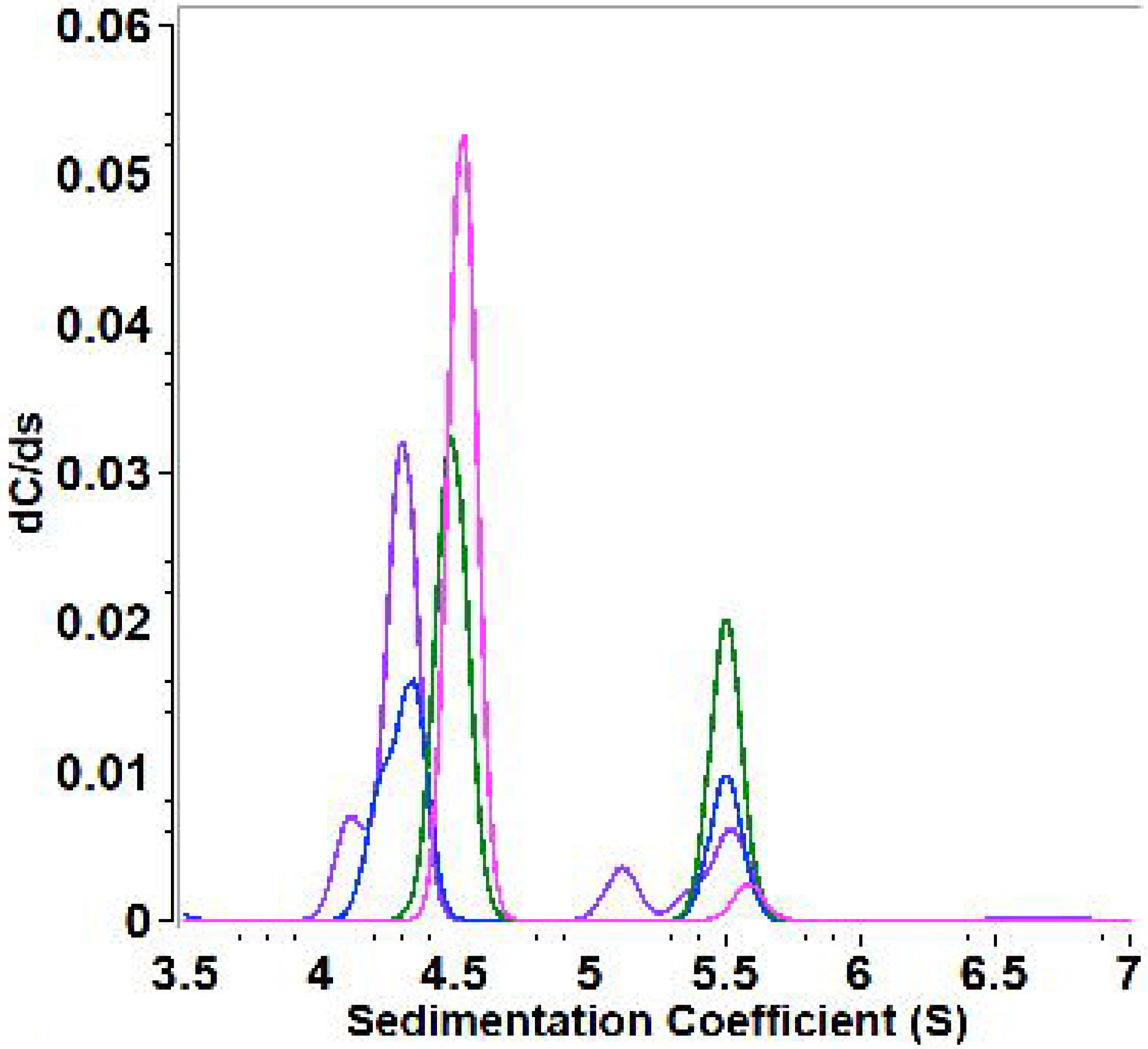
Homogeneity studies of MVEV and PowV non-coding terminal regions using SV-AUC, where x-axis represents sedimentation coefficient distribution. The SV peaks at ∼4.5 S for each RNA represent monomeric fractions. Colour scheme is identical to Figure 2.

### 3.3. Low-resolution structural studies of RNAs

Small-angle X-ray scattering allows solution structure studies of biomolecules and their complexes, albeit at low-resolution. Recent developments with instrumentation in terms of an HPLC unit connected inline with SAXS detection allows for data collection of monodispersed sample devoid of aggregated and degraded products [51,52]. As outlined in the Materials and Methods section, we collected selected HPLC-SAXS data for all four RNAs, followed by the selection of data from a monodispersed peak, buffer-subtraction, and merging of selected datasets. The merged SEC-SAXS data are presented in Figure 4A. Subsequently, the merged data were processed using the Guinier method (plot of (*I*(*q*)) vs. (*q*^2^)), which aids detection of purity and allows determination of the *R*_*g*_ from the data belonging to the low-*q* region [38]. Figure 4B presents the Guinier plots for 5’ and 3’ TRs of MVEV and PowV, where the linearity for low-*q* data demonstrates that indeed all four RNAs are monodispersed and devoid of any aggregation. Based on the Guinier analysis, we obtained *R*_*g*_ values of 29.01 ± 0.22 Å, 45.72 ± 0.48 Å, 34.65 ± 0.19 Å and 35.84 ± 0.13 Å for MVEV 5’ TR, MVEV 3’ TR, PowV 5’ TR and PowV 3’ TR respectively (see Table 1 for more details). Once we confirmed the monodispersity, we processed the raw SAXS data from Figure 4A to obtain dimensionless Kratky plots (*I(q)/I(0)*(q*R*_*g*_)^*2*^ *vs q*R*_*g*_) that allow detection of the folding state of biomolecules [40,52]; for example, globular-shaped biomolecules in solution are observed with a well-defined maximum value of 1.1 at *q*\**R*_*g*_ =√3 [39]. However, the dimensionless Kratky plots for 5’ and 3’ TR of MVEV and PowV under investigation demonstrate that all the samples are well-folded and extended in solution (Figure 4C).

**Figure 4.**
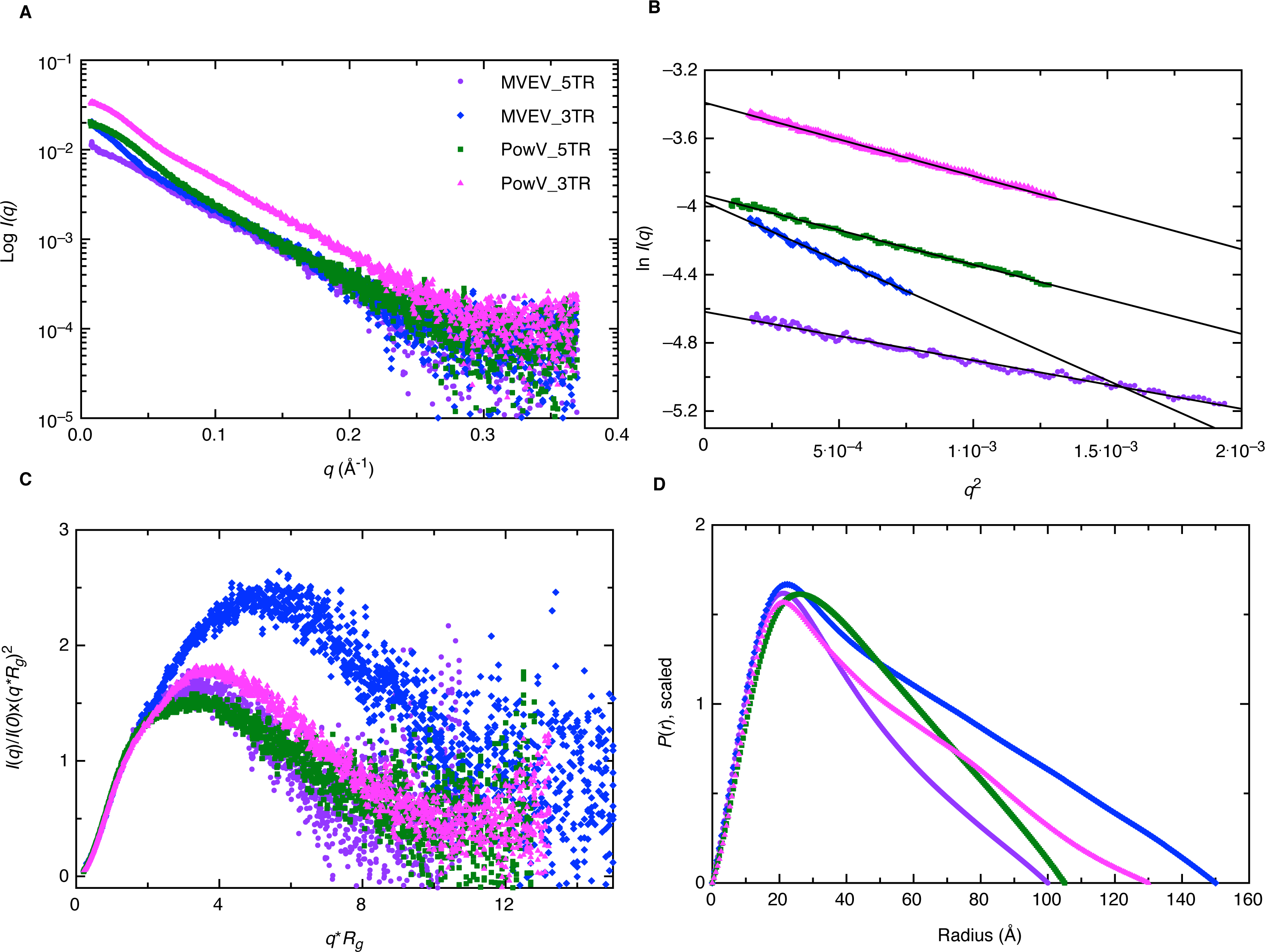
Characterization of MVEV and PowV terminal regions using SAXS. (A) A plot of scattering intensity (Log I(q)) versus scattering angle (q = 4πsinθ/λ) representing merged SAXS data for MVEV and PowV. (B) Guinier plots (plot of ln(I(q)) versus q^2^) representing the homogeneity of samples and allowing determination of R_g_ based on the low-angle region data. (C) Dimensionless Kratky plots (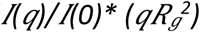 versus q.R_g_) for all four RNA samples demonstrating their extended structures. (D) Pair-distance distribution (P(r)) plots for all four RNA samples representing their maximal particle dimensions and allowing the determination of Rg from the entire SAXS dataset.

Next, using program GNOM, we performed an indirect Fourier transformation to convert the reciprocal-space information (ln(*I*(*q*)) vs. (*q*), Figure 4A) into the real space electron pair-distance distribution function (*P*(*r*), Figure 4D)) to obtain *R*_*g*_ and *D*_*max*_ for all four RNAs. Note that unlike Guinier analysis, which is restricted to the data in the low-*q* region, the *P*(*r*) analysis utilizes a wider-range of the dataset and aids reliable determination of *R*_*g*_ as well as of *D*_*max*_. As outlined in Table 1, based on *P*(*r*) analysis, we obtained the *D*_*max*_ of 100 Å, 150 Å, 105 Å, and 130 Å for MVEV 5’ TR, MVEV 3’ TR, PowV 5’ TR, and PowV 3’ TR respectively. Furthermore, the *R*_*g*_ values of 30.16 ± 0.11 Å, 46.16 ± 0.18 Å, 34.48 ± 0.06 Å and 38.39 ± 0.08 Å for MVEV 5’ TR, MVEV 3’ TR, PowV 5’ TR and PowV 3’ TR respectively were obtained. These values are very similar to those obtained from Guinier analysis indicating that the data are suitable for low-resolution shape reconstruction. The shape of the *P*(*r*) plot is indicative of the solution conformation of biomolecules. For example, for a globular-shaped protein, we would have expected a bell-shaped *P*(*r*) distribution curve with a maximum at ∼*D*_*max*_/2 [53]. However, as presented in Figure 4D, all four RNAs display skewed bell-shaped curves with extended tails, suggesting their extended structures in solution.

To obtain a low-resolution solution structure of each RNA, we employed program DAMMIN [42] that utilizes simulated-annealing protocol and allows the incorporation of *P*(*r*) data (*i.e. D*_*max*_ and *R*_*g*_ values as constraints). We calculated a total of 12 models for 5’ and 3’ TRs of MVEV and PowV and noted that individual models had an excellent agreement between the experimentally obtained scattering data and the calculated scattering data, as the χ^2^ values in each case are ∼ 0.8 (see Table 1). Next, we employed program DAMAVER to rotate, align and obtain an averaged filtered structure for each RNA [44]. In each case, the goodness of the superimposition of individual models was estimated by the overlap function—normalized spatial discrepancy (NSD). As presented in Table 1, the low NSD values suggest that 12 models in each case are highly similar to each other. The averaged filtered structures for the 5’ and 3’ TRs of MVEV and PowV are presented in Figure 5, which indicates that indeed, overall these RNAs have extended structures in solution.

**Figure 5.**
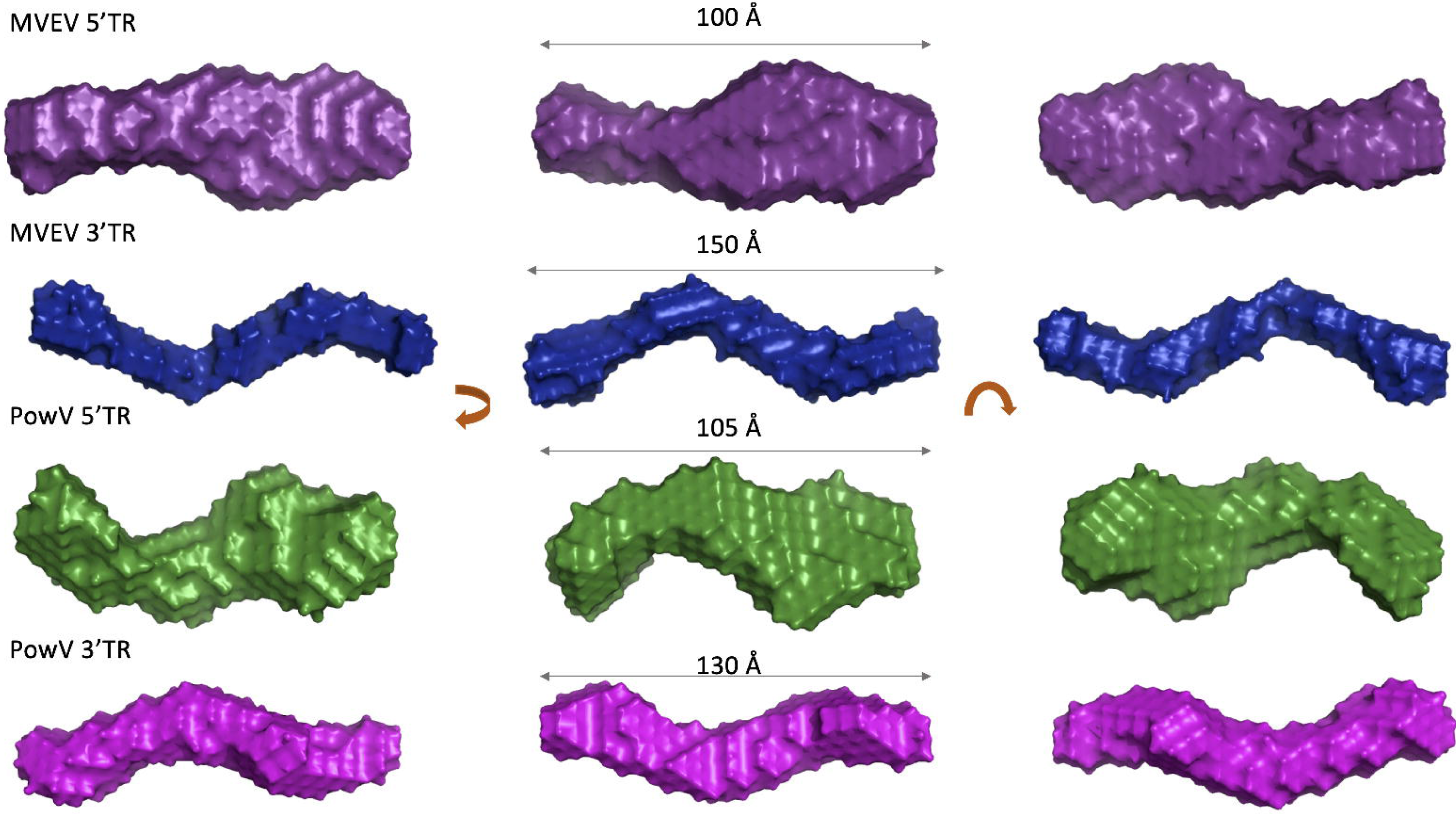
Determination of low-resolution structures using SAXS, which indicates that all RNA molecules adopt an extended structure in solution. For each terminal region RNA, the left and right panels represent 180º rotated view on the x-axis and y-axis, respective of the view presented in the middle panel. Dimensions represent the D_max_.

### 3.4. Computational modeling of RNA structures

SAXS is an outstanding method that allows the determination of solution structures of biomolecules, albeit at low-resolution. Techniques such as X-ray crystallography and nuclear magnetic resonance spectroscopy, provide high-resolution structural information but have limited applications for RNA structural studies due to challenges obtaining high-quality crystals and RNA-labeling, respectively. An alternative to this approach is to employ computational modeling of RNA structures and screen those structures using experimental data. We employed the MC-Fold/MC-Sym pipeline developed by Parisien and Major [47]. For each RNA, we calculated 100 structures using the MC-Sym pipeline based on the secondary structure predicted using MC-Fold (Figure 1). Next, we used program CRYSOL [48] to calculate X-ray scattering profiles of 100 structures of each RNA – 5’ and 3’ TRs of MVEV and PowV. CRYSOL also allows the determination of *R*_*g*_ from these structures. We also calculated a χ^2^ value between the X-ray scattering data of MC-Sym derived structures that were calculated using CRYSOL and experimentally collected X-ray scattering data. Figure 6 presents the *R*_*g*_ (*y*-axis left-hand side, solid black circles) and χ^2^ values (*y*-axis right-hand side, grey double circles) for each RNA system suggesting that the RNA molecules under investigation can adopt a variety of conformations. For example, in the case of 5’ TR of MVEV, the distribution of *R*_*g*_ values rage from 30 ± 5 Å, whereas the experimentally determined *R*_*g*_ for this construct is 30.16 Å. Similarly, for PowV 3’ TR, experimentally determined *R*_*g*_ is 38.39 Å, which largrly alings with the distribution of *R*_*g*_ values of MC-Sym derived structures. On the other hand, the MC-Sym derived models for MVEV 3’ TR mainly under-represent the *R*_*g*_ values when compared to experimental *R*_*g*_ of 46.16 Å. In contrast, for PowV 5’TR, Mc-Sym derived structures have a higher *R*_*g*_ distribution (∼45Å) compared to the experimentally determined *R*_*g*_ (34.48 Å). Moreover, as presented in Figure 6, the χ^2^ values for MC-Sym derived 3’ TR of MVEV and PowV structures have better distribution around 1.5 χ^2^, compared to MVEV and PowV 5’ TRs.

**Figure 6.**
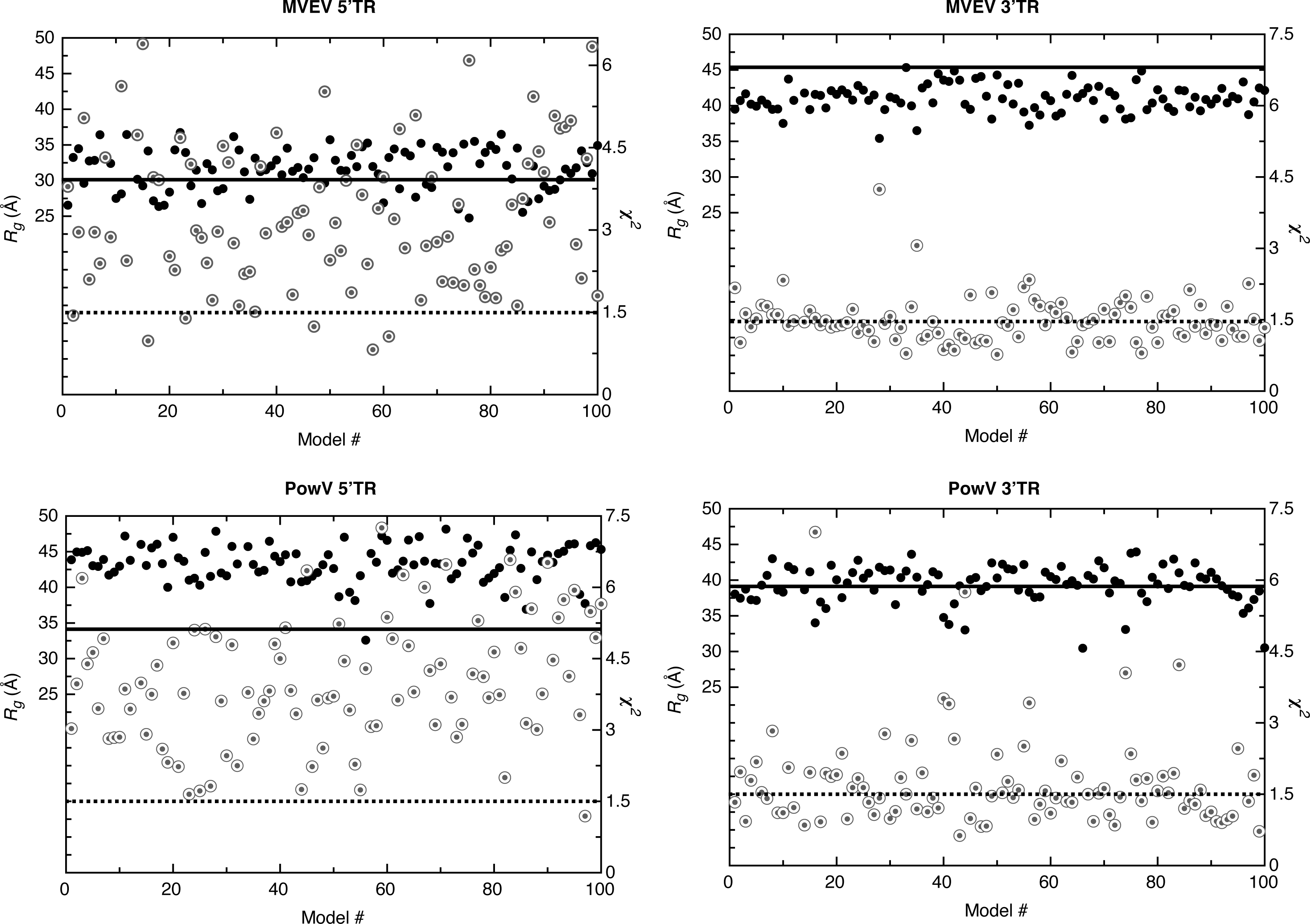
Screening of high-resolution structures calculated using MC-Sym [47] for MVEV 5’ TR, MVEV 3’TR, PowV 5’TR and PowV 3’TR. For each RNA, the x-axis represents model numbers (total 100 models), whereas the y1-axis (solid black circles) and y2-axis (grey double circles) represent R_g_ (Å) and χ^2^ values, respectively, calculated using CRYSOL package [48]. For each plot, the solid dark line corresponds to y1-axis and represents the value for experimentally determined R_g_ (Å) as presented in Table 1. The dotted line corresponds to y2-axis and represents a χ^2^ value of 1.5. These plots indicate that MC-Sym derived structures represent a wide-range of conformations these RNAs can theoretically adopt.

### 3.5 Combination of Computational Modeling and Experimental Low-Resolution SAXS Structures

We selected ∼10 MC-Sym derived structures with the lowest χ^2^ values to further investigate if they align with the low-resolution averaged filtered models for each RNA system. Figures 7 and 8 present four of the best χ^2^ fit MC-Sym derived structures for MVEV and PowV, indicating that these structures agree well with the low-resolution models based on SAXS data. Overall, MC-Sym derived structures for the 3’ TRs of MVEV and PowV (Figure 8) indicate that they adopt highly extended structures mainly in the double-stranded regions. In contrast, the 5’ TRs of MVEV and PowV are more dynamic with less double-stranded regions (Figure 7). Furthermore, these results indicate that the 5’ TR structures have a higher content of un-paired/loop regions compared to the 3’ TRs which contains mainly double-stranded regions.

**Figure 7.**
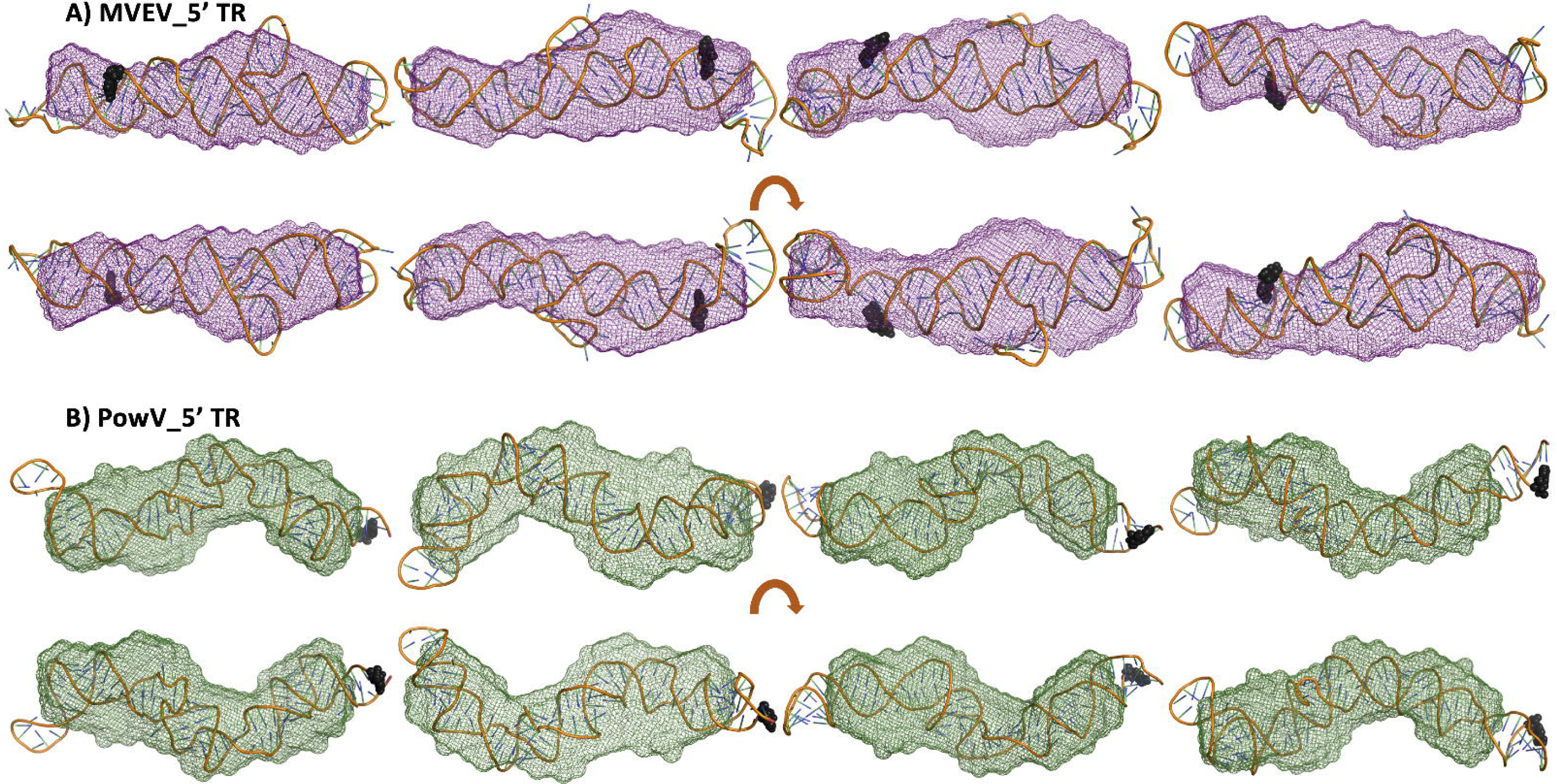
High-resolution structures calculated using MC-SYM overlaid with low-resolution SAXS models. A) MVEV 5TR B) and PowV 5TR. Bottom panels in both cases represent a 180° rotation about the x-axis represented in the top panels. Black spheres represent 5’ terminal region on each construct.

**Figure 8.**
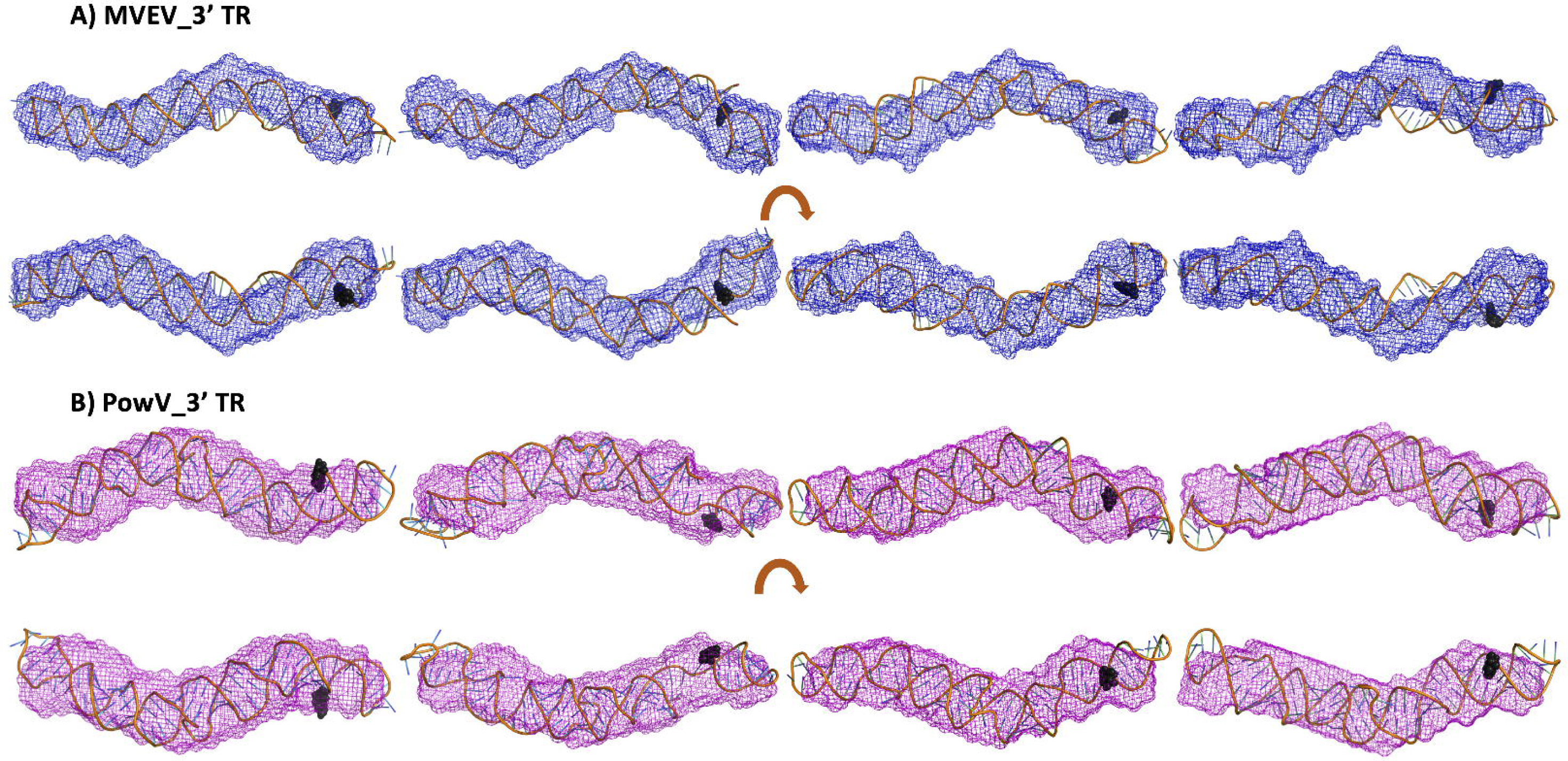
High-resolution structures calculated using MC-SYM overlaid with low-resolution SAXS models. A) MVEV 3TR B) and PowV 3TR. Bottom panels in both cases represent an 180° rotation about the x-axis represented in the top panels. Black spheres represent 5’ terminal region on each constructs.

## 4. Discussion

MVEV is endemic to northern Australia and has seen an increase in prevalence in the last decade. With extensive human development in areas affected, the likelihood of an outbreak is only increasing. Similarly, PowV is an under-studied flavivirus despite its increasing incidence and severe neurological effects. Patients infected with almost any flaviviruses, including PowV and MVEV, are only treated with symptomatic support. Treatment of the viral infection is nonexistent, resulting in a necessity for further investigation into these potentially deadly viruses. Therefore, further work is required to gain detailed insights into the viral life cycle to better prepare ourselves for potential future outbreaks.

Lowest energy secondary structure predictions (Figure 1) reveal that each RNA molecule varies slightly due to increasing energy, with the exception of PowV 5’ TR, and to a lesser degree MVEV 3’ TR. The structure with the lowest predicted energy of PowV 5’ TR displays a large change in secondary structure with only a small change in energy (−103.07 vs −102.43). The secondary structure information for MVEV 3’ TR indicates that an increase in energy could result in the formation of a second stem-loop with a small energy change (−104.44 to −103.29). This suggests that the RNA molecules can adopts multiple different conformations in solution.

We prepared all four RNAs using *in-vitro* transcription with T7 polymerase and purified using SEC (Figure 2B), which indicates that these RNAs are able to form dimer/oligomers in solution. The fractions indicating a homogenous preparation were pooled together and analyzed using Urea-PAGE (Figure 2B), where all four RNAs displayed a single band, indicating the absence of degradation. The resolution of the denaturing gel was not enough to discern the difference in molecular weights between MVEV (∼32 kDa) and PowV (∼37kDa) RNA constructs (Table 1). Therefore, we employed sedimentation velocity experiments using analytical ultracentrifuge which determines the composition and anisotropy of biomolecules in solution [50,54,55]. Figure 3 presents the sedimentation coefficient distribution of SEC-purified RNA preparation. As evident by the peaks between 5S and 6S, all these RNAs can form higher-order oligomers to a small degree, despite removing many of them by means of SEC purification. This is consistent with the HPLC data from SAXS (not shown) which also shows a minor amount of oligomerization for each RNA. Another explanation for these minor oligomerization peaks could be the RNA adopting different conformations, as was seen in the predicted secondary structure(s) of PowV 5’ TR (Figure 1) where a small energy change was seen to cause the RNA to adopt a different conformation. This prediction could explain the increased peak that is evident at 5.5 S (Figure 3), for both PowV 5’ TR and MVEV 3’ TR. Additionally, the peak at ∼ 4 S for MVEV 5’ TR indicates minor degradation. More importantly, the differences in RNA size and shape are indicated by their sedimentation coefficient values (Table 1). For example, both the TRs of MVEV have a *M*_*w*_ of ∼ 32 kDa and produce very similar S values (4.27 S and 4.3 S, for 5’ and 3’ TR, respectively). Furthermore, the S values for 5’ and 3’ TRs of PowV are 4.50 S and 4.53 S, this similarity correlates with the similarities of their *M*_*w*_ of ∼ 37 kDa (Table 1).

Small-angle X-ray scattering (SAXS) has emerged as an excellent complimentary structural-biophysical method which aids solution structural studies of RNA, proteins, and their complexes, albeit at low-resolution [40,43,51,56-59]. Therefore, solution X-ray scattering is employed for biological systems where obtaining high-quality crystals or labeling of biomolecules that presents challenges [21,43,45,57,58,60]. In this study, we employed an HPLC-SAXS set-up to collect scattering data from a preparation free of any aggregation or degradation. The monodispersed preparation was confirmed by Guinier analysis and shows excellent linearity of fit in the low-*q* region (Figure 4B). Furthermore, Guinier analysis also provided *R*_*g*_ of all four RNAs (based on the low-*q* region), which are highly similar to those calculated by means of *P*(*r*) analysis (Figure 4D, Table 1). We also performed dimensionless Kratky analysis (Figure 4C) that demonstrates each RNA adopts an elongated structure, with a low amount of flexibility. The *P*(*r*) distribution reveals a quick increase to the highest point, and then a gradual decrease where the electron-pair-distance approaches zero. A globular molecule would result in a gaussian-like distribution, which is not evident in any of the RNA, suggesting their elongated nature. The most extended molecule is MVEV 3’ TR resulting in a *D*_*max*_ of 150 Å, even though its molecular weight is 31.57kDa (Table 1), lower than both 5’ and 3’ TR of PowV (Table 1). Low-resolution structure modeling (Figure 5) further confirms that MVEV 3’ TR adopts a much thinner and elongated structure, suggesting that this RNA is almost entirely helical. This is also evident for PowV 3’ TR which also displays an extended conformation. This elongation is consistent with the predicted secondary structure of flaviviral 3’ terminal regions where the last ∼100 nucleotides are suggested to almost entirely base pair into a large system [21,61-63]. Compared to the 3’ TRs, low-resolution structures of 5’ TRs of MVEV and PowV are less elongated. However, their low-resolution structures appear to be consistent with the predicted secondary structures of flaviviral 5’ TRs, indicating that the 5’ TRs adopt more dynamic structures [25,61-63]. Overall, MVEV and PowV TR structures are consistent with previously published low-resolution structures of flaviviral RNA such as the Zika, and West Nile virus [21,25,64].

SAXS can be combined with high-resolution structures/homology models of individual domains as well as with computational studies, to compliment predicted structures [21,40,57,60,65]. RNA structures of high resolution are hard to determine through crystallization and labeling studies mentioned above; therefore, an attractive alternative to calculating high-resolution structures is computational modeling. To this end, we employed the MC-Fold/MC-SYM pipeline to reconstruct all-atom structures and screen those using experimentally collected X-ray scattering data to identify structures that are likely adopted by these RNAs. Using CRYSOL [48], we calculated *R*_*g*_ for each structure for each TR, as well as back-calculated X-ray scattering data, and aligned them with experimentally collected scattering data (Figure 6). This analysis demonstrated that MVEV 3’ TR and PowV 3’ TR exhibit a strong correlation between the experimentally determined *R*_*g*_ and the calculated *R*_*g*_ from MC-Sym-derived structures. Similarly, for both of these RNAs, *χ*^*2*^ values indicate agreement between scattering data that were experimentally collected and calculated from MC-Sym-derived structures, where the *χ*^*2*^ values were close to ∼1.5 for most structures (Figure 6). On the other hand, the *χ*^*2*^ values for MC-Sym-derived structures for 5’ TRs of MVEV and PowV have a much larger distribution (from ∼1.5 to ∼6). A similar trend was also observed for *R*_*g*_ for both 5’ TRs where the MC-Sym derived structures displayed a wider distribution of *Rg*, suggesting that these RNAs could adopt multiple conformations. On the other hand, the correlation between *R*_*g*_ and *χ*^*2*^ was far more pronounced in the more elongated TR RNAs - PowV 3’ TR and MVEV 3’ TR (Figures 5 and 6). Finally, we selected the ∼10 MC-Sym-derived structures, which had the lowest *χ*^*2*^ values when compared to experimental data and aligned them into the low-resolution SAXS envelopes. Selected groups of structures are presented in Figures 7 and 8, which indicate the computationally calculated structures agree well with the experimental data, especially for 3’ TRs. For the 5’ TRs, we observe dynamic structures; however, they maintain overall, similar conformation in solution. Furthermore, the bulge regions in both 5’ TRs also agree well with their secondary structures, suggesting that these secondary structures could be conserved and do not undergo conformational changes. An added benefit of these computationally derived structures is the addition of directionality of the RNAs (the black spheres represent the 5’ end of each molecule in Figures 7 and 8), which is otherwise very challenging to decipher unless the molecule of interest is altered [66].

## 5. Conclusions

In this study, we have *in-vitro* transcribed MVEV and PowV terminal RNA regions and performed their native purification, studied their homogeneity using analytical ultracentrifugation, determined their low-resolution structures and combined computational modeling to obtain high-resolution structural models. We demonstrated that the combination of SAXS and computational methods allow further in-sights into structural details of RNA molecules.

## Author Contributions

Project was conceptualized by TM and TRP, who also performed SAXS data collection, and data analysis. TM prepared RNA samples. AH and TM collected AUC data. AH, TM and BD processed AUC data. All authors contributed in manuscript writing and figures preparation.

